# Delivery of GalNAc-conjugated splice-switching ASOs to non-hepatic cells through ectopic expression of asialoglycoprotein receptor

**DOI:** 10.1101/382606

**Authors:** Juergen Scharner, Sabrina Qi, Frank Rigo, C. Frank Bennett, Adrian R. Krainer

## Abstract

Splice-switching antisense oligonucleotides (ASOs) are promising therapeutic tools to target various genetic diseases, including cancer. However, *in vivo* delivery of ASOs to orthotopic tumors in cancer mouse models or to certain target tissues remains challenging. A viable solution already in use is receptor-mediated uptake of ASOs via tissue-specific receptors. For example, the asialoglycoprotein receptor (ASGP-R) is exclusively expressed in hepatocytes. Triantennary GalNAc (GN3)-conjugated ASOs bind to the receptor and are efficiently internalized by endocytosis, enhancing ASO potency in the liver. Here we explore the use of GalNAc-mediated targeting to deliver therapeutic splice-switching ASOs to cancer cells that ectopically express ASGP-R, both *in vitro* and in tumor mouse models. We found that ectopic expression of the major isoform ASGP-R1 H1a is sufficient to promote uptake and increase GN3-ASO potency to various degrees in all tested cancer cells. We show that cell-type specific glycosylation of the receptor does not affect its activity. *In vivo*, GN3-conjugated ASOs specifically target subcutaneous xenograft tumors that ectopically express ASGP-R1, and modulate splicing significantly more strongly than unconjugated ASOs. Our work shows that GN3-targeting is a useful tool for proof-of-principle studies in orthotopic cancer models, until endogenous receptors are identified and exploited for efficiently targeting cancer cells.

## Introduction

Antisense oligonucleotides (ASOs) have become an important therapeutic tool that has vastly expanded the types of diseases that can be treated in a targeted and safe way.^1^ ASOs are short, chemically modified, single-stranded oligonucleotides that bind to specific complementary RNA sequences by Watson-Crick base pairing. Based on their design, they either elicit degradation of the RNA target by RNAse H-mediated cleavage, or act as steric blockers of protein or RNP binding, e.g., to modulate RNA splicing and induce an isoform switch or restore protein expression.^2-4^ ASOs have also successfully been deployed to neutralize micro-RNAs,^5^ increase translation,^6^ and to reduce nonsense-mediated mRNA decay.^7^ Currently, a major limitation of the use of ASOs is their efficient delivery to the intended target tissue.^8^ ASOs delivered systemically *in vivo* are cleared rapidly from the blood and primarily accumulate in the liver and kidney, although ASO presence and activity have been detected in a wide variety of tissues.^9, 10^ Once ASOs enter cells, they have very long half-lives, ranging from 2-4 weeks in the liver^10^ to 4-6 months in the CNS.^11, 12^ Antisense drugs already approved by the FDA include those designed against diseases affecting tissues that are either self-contained or easy to target, such as eye, liver and CNS.^1^ An ASO targeting skeletal muscle has also been conditionally approved by the FDA for Duchenne muscular dystrophy, though its efficacy is limited by inefficient muscle uptake.^13^ There are extensive ongoing efforts to develop methods for efficient, tissue-specific targeting, including aptamers, lipid nanoparticles, cell-penetrating peptides, antibodies, and receptor ligands.^8^ Tissue-specific targeting is especially crucial for cancer therapies, as ASOs are diluted out in rapidly dividing cells, thus requiring higher and more frequent dosing, compared to post-mitotic tissues.^14, 15^

A well-established receptor/ligand system to target hepatocytes already in use in clinical trials is the asialoglycoprotein receptor (ASGP-R).^16^ ASGP-Rs are primarily expressed in hepatocytes, and play an important role in clearing glycoproteins from the blood through clathrin-mediated endocytosis. There are five receptor isoforms encoded by two different genes, *ASGR1* and *ASGR2*. Two isoforms lack the transmembrane domain and are soluble; the remaining three isoforms (ASGP-R1 H1a, ASGP-R H2b and H2c) are membrane-bound, and homo- and hetero-oligomerize upon ligand binding on the cell surface, before endocytosis. The ligand specificity is determined by the receptor-oligomer composition.^17^ ASOs conjugated with N-acetyl galactosamine, a natural ASGP-R ligand, are efficiently bound by the receptor and internalized in hepatocytes; as a result, GN3-conjugated ASOs increase the potency of liver-targeting ASOs *in vivo* by ~10-fold.^18^ Cancer-specific receptors, such as the IL-13Rα2 or EGFRvIII receptors, which are specifically expressed or amplified glioblastomas, are already being tested for targeted therapies using ligand and aptamers, but are not yet widely available.^19-21^

Here we aimed to adopt the hepatic ASGP-R/GN3 receptor/ligand system for targeted delivery of GN3-conjugated ASOs to non-hepatic cancer cell lines, by ectopically expressing ASGP-R. Early work characterizing receptors in mouse fibroblasts, and more recent work in HEK293T cells, showed that ASGP-R is functional when expressed ectopically.^22, 23^ Furthermore, ASGP-R expression can enhance the potency of unconjugated ASOs *in vitro* and *in vivo*, likely via direct interaction between the P=S-modified ASO backbone and the receptor.^23, 24^ Ectopic expression could therefore be used for proof-of-principle experiments to test the efficacy therapeutic ASOs, both *in vitro* and *in vivo*, until endogenous, cancer/tissue-specific receptors are identified. We expressed ASGP-R isoforms in various human cancer cell lines to identify cell-type specific differences in receptor activity, as well as culture conditions that influence cellular ASO uptake. We found that ectopic expression of the major isoform ASGP-R1 H1a is sufficient to promote uptake and increase GN3-ASO potency in some, but not all cancer cells we tested. *In vivo*, GN3-conjugated ASOs specifically targeted subcutaneous xenograft tumors expressing ASGP-R1, and induced significantly stronger splice-modulation than unconjugated ASOs. We conclude that GN3-targeting is a useful method to test therapeutic ASOs in proof-of-principle *in vivo* studies employing orthotopic cancer models.

## Results

### ASGP-R promotes GN3-conjugated ASO uptake and efficacy in U87 cells

GN3-conjugated oligonucleotides (siRNAs and gapmer ASOs) have been successfully used to target hepatocytes *in vivo* via ASGP-R mediated endocytosis. There is extensive effort in the field to identify new receptors, with the aim to deliver ligand-conjugated ASOs to other target tissues or tumor cells. While comparable receptor/ligand systems are being developed for other tissues, we aimed to test whether ectopic expression of ASGP-R in non-hepatic cells can promote uptake and efficacy of GN3-conjugated splice-modulating ASOs, for proof-of-principle experiments *in vitro* and *in vivo*.

Hepatocytes express multiple ASGP-R isoforms encoded by two genes (*ASGR1* and *ASGR2*), and these isoforms oligomerize on the cell surface upon ligand binding. Because it was unclear which isoforms are required to induce GN3 binding and endocytosis, we cloned all membrane-bound ASGP-Rs into retroviral and lentiviral expression vectors, to test them in non-hepatic cells (Fig. 1A). HepG2 hepatocellular carcinoma cells, which served as a control, express low levels of both ASGP-R1 and ASGP-R2 endogenously (Fig. 1B). In U87 glioblastoma cells, ASGP-R is not expressed. Western blots show strong expression of isoform H1a only in infected cells (Fig.1B). ASGR2 isoforms are retained in the ER and rapidly degraded when expressed alone in HEK293 cells.^22, 23^ ASGP-R2 isoforms expressed individually in U87 cells were not stable, and required the presence of isoform H1a for stability and proper localization, which is consistent with the literature (Fig. 1B-C). We confirmed this observation by immunostaining, which showed accumulation of H2b near the nucleus (consistent with ER localization) when expressed alone (Fig. 1C, arrow heads).

**Figure 1.**
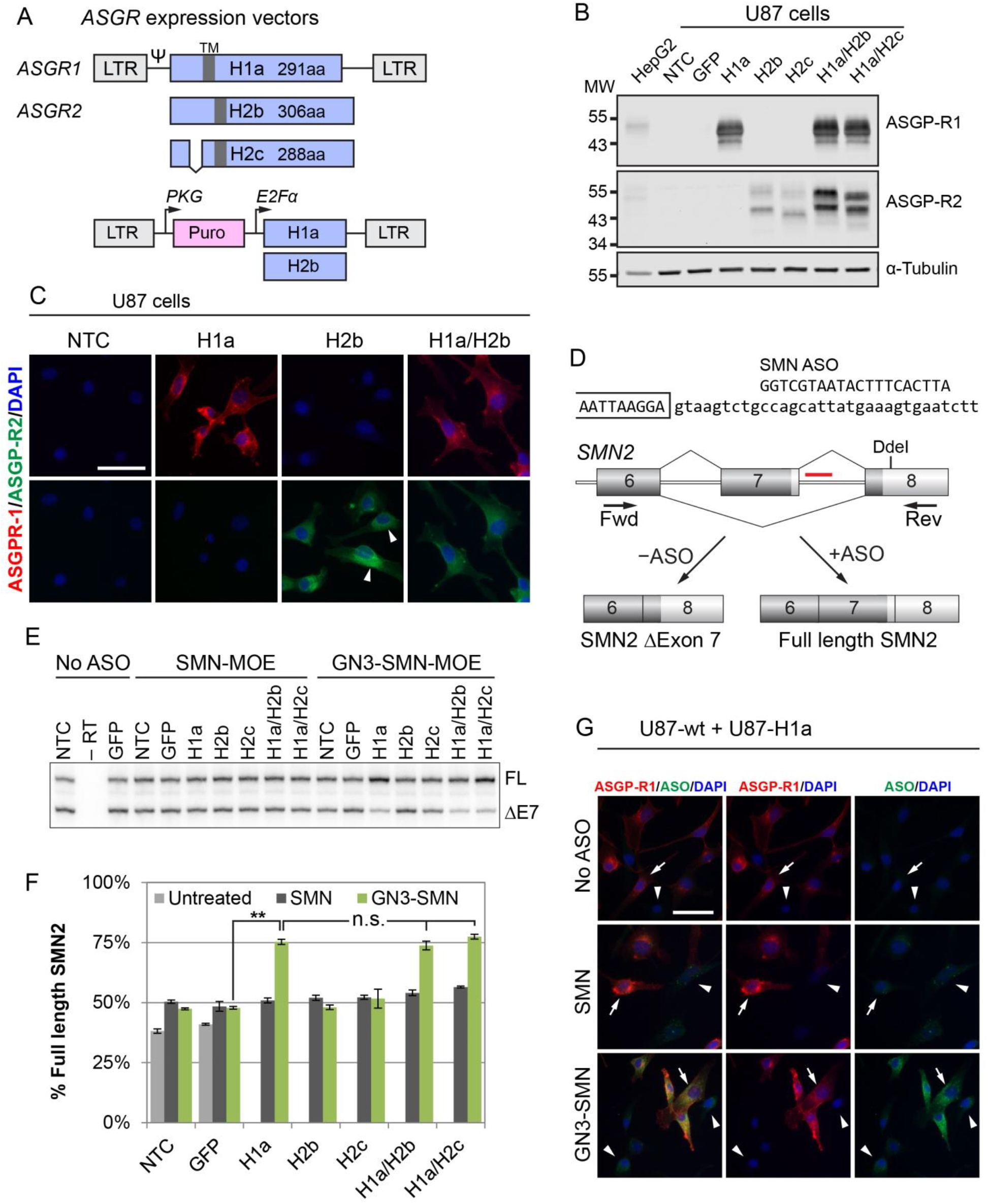
Ectopic expression of ASGP-R1 in U87 cells increases efficacy of GN3-SMN-ASO *in vitro*. (A) Major and minor transmembrane domain (TM)-containing ASGP-R isoform cDNAs were cloned into retroviral and lentiviral expression vectors, as indicated. The lentiviral vector carries the selection marker puromycin. ASGP-R H2c has a short 18-amino-acid deletion in the intracellular domain of the receptor. (B) Western blot confirms expression of major and minor ASGP-R isoforms in U87 glioblastoma cells. The expression level of H1a is approx. 6-fold higher than the endogenous expression level in HepG2 cells, normalized to tubulin. H2a and H2b isoforms are detectable when expressed alone, but are stabilized in the presence of H1a. (C) Micrographs of U87 cells expressing ASGPR-H1a and H2b alone, and in combination. Cells were stained for ASGP-R1 (red), ASGP-R2 (green), and DAPI (blue). Arrowheads indicate non-uniform distribution of ASGP-R H2b, consistent with ER localization. Scale bar, 50 μm. (D) The SMN ASOs used in this study bind to intron 7 of *SMN2* and promote exon 7 inclusion. Full-length *SMN2* mRNA was quantified by radioactive RT-PCR; the product was digested with DdeI to separate *SMN1* from *SMN2* products. (E) U87 cells expressing major and minor ASGP-R isoforms alone or in combination were incubated with 300 nM unconjugated (SMN-MOE) or GalNAc-conjugated SMN-MOE ASOs (GN3-SMN-MOE) for 5 d by free-uptake. Representative radiograph shows full-length *SMN2* (top band) and *SMN2* Δ exon 7 (bottom band). (F) Quantification of full-length *SMN2* in ASO-treated U87 cells. N=3 independent retroviral transductions; bar graphs represent mean +/- S.E.; ** *P* < 0.01; NTC, no treatment control; n.s., not significant. (G) U87 and U87-H1a cells exposed to unconjugated and GN3-conjugated SMN-ASOs for 24 hrs. Cells were stained for ASGP-R1 (red), ASO (green), and DAPI (blue). Arrows indicate ASGP-R1-expressing U87 cells, and arrowheads indicate ASGP-R1-negative cells. Scale bar, 50 μm.

To test whether ASO-uptake and efficacy is improved in ASGP-R-expressing U87 cells, we used 20mer unconjugated (SMN-MOE) and GN3-conjugated (GN3-SMN-MOE) fully MOE-PS modified splice-modulating ASOs. The ASO moiety binds to an intronic splice silencer site in intron 7 of *SMN2*, blocking hnRNPA1 from binding and thereby promoting exon 7 inclusion.^25^ We measured *SMN2* exon 7 inclusion by radioactive RT-PCR with primers located in exons 6 and 8 (Fig. 1D). We delivered ASOs to control and receptor-expressing U87 cells by free uptake, in the absence of any transfection reagent, for 5 days at a final concentration of 300 nM. Untreated wild-type and control U87 cells (NTC, GFP) expressed 38.1% (±1.5%) and 41.0% (±0.6%) full-length *SMN2*, respectively (Fig. 1E-F). The proportion of full-length *SMN2* mRNA significantly increased in wild-type U87 cells exposed to unconjugated and GN3-conjugated SMN-ASOs, to a similar extent (50.4±0.7% and 47.4±0.4% respectively, Fig. 1F). In U87 cells expressing ASGP-R isoforms H1a, H2b and H2c alone or in combination, unconjugated SMN-ASOs induced exon 7 inclusion to a similar extent as in ASO-treated control cells. GN3-conjugated ASOs, in contrast, performed significantly better in ASGP-R1-expressing cells. U87-H1a cells expressed 50.9±1.0% FL-SMN2 when treated with unconjugated SMN-ASOs, and 75.3±1.1% FL-SMN2 when treated with GN3-conjugated SMN-ASOs. In cells expressing two ASGP-R isoforms (U87-H1a/H2b and U87-H1a/H2c), the expression of FL-SMN2 was not significantly different from that in U87 cells expressing the H1a isoform alone. This observation suggests that in U87 cells, expression of the ASGP-R H1a isoform is sufficient to induce uptake of GN3-conjugated ASOs. We also tested cells expressing all three membrane-bound isoforms, and saw no statistically significant improvement over U87-H1a cells (Fig. S1).

We confirmed specific GN3-ASO targeting to U87-H1a cells by immunohistochemistry. We exposed a mixture of U87-wt and U87-H1a cells to 300 nM unconjugated or GN3-conjugated ASO for 24 hrs, and stained for ASGP-R and ASO, the latter with an antibody that recognizes the PS backbone. As shown in Fig. 1G, internalization of unconjugated ASOs was inefficient in both U87-wt and U87-H1a cells. In contrast, GN3-conjugated ASO-staining was much stronger in U87-H1a cells than in U87-wt cells, demonstrating greatly enhanced uptake efficiency. However, Gn3-ASO staining was detectable in U87-wt cells, showing that GN3-ASO can be internalized through alternative pathways, or the GN3-ligand was metabolized releasing the ASO over the incubation period.

### GN3-ASO uptake efficiency correlates with ASGP-R1 expression level

Asialoglycoprotein receptors are among the most highly expressed receptors, though there is evidence that only a small pool of receptors is required for efficient ASO internalization.^23^ To test whether the expression level positively influences GN3-ASO activity, we transiently expressed ASGP-R1 in U87 cells by transduction with increasing retroviral titers (U87-low/medium/high, Fig. 2A), and then treated these cells with 0-30 μM SMN-ASOs in 7-point dose-response experiments for 5 days. Fig. 2B and C show that the proportion of FL-SMN2 increased with the ASO dose in U87-wt cells. Unconjugated and GN3-conjugated ASOs had very similar effects in cells lacking ASGP-R1. However, with increasing levels of ASGP-R1, the potency of GN3-conjugated ASOs (green curve) increased, compared to unconjugated ASOs (black curve). In U87-high cells, the GN3-ligand increased the ASO potency by approximately 50-fold, compared to its unconjugated counterpart. In U87-low and U87-medium cells, the potency of the GN3-conjugated ASO increased only approximately 10-fold and 20-fold, respectively (Fig. 2C). Interestingly, transient expression of ASGP-R1 in A172 cells, a different human glioblastoma cell line, only resulted in a 3-fold increase in potency of the GN3-conjugated ASO, even in cells expressing high levels of ASGP-R1 (Fig. S2). This poor performance of ASGP-R1 in A172 cells, despite high levels of expression of ASGP-R1 suggests that expression of the receptor is insufficient to enhance activity of the GN3-conjugated ASO.

**Figure 2.**
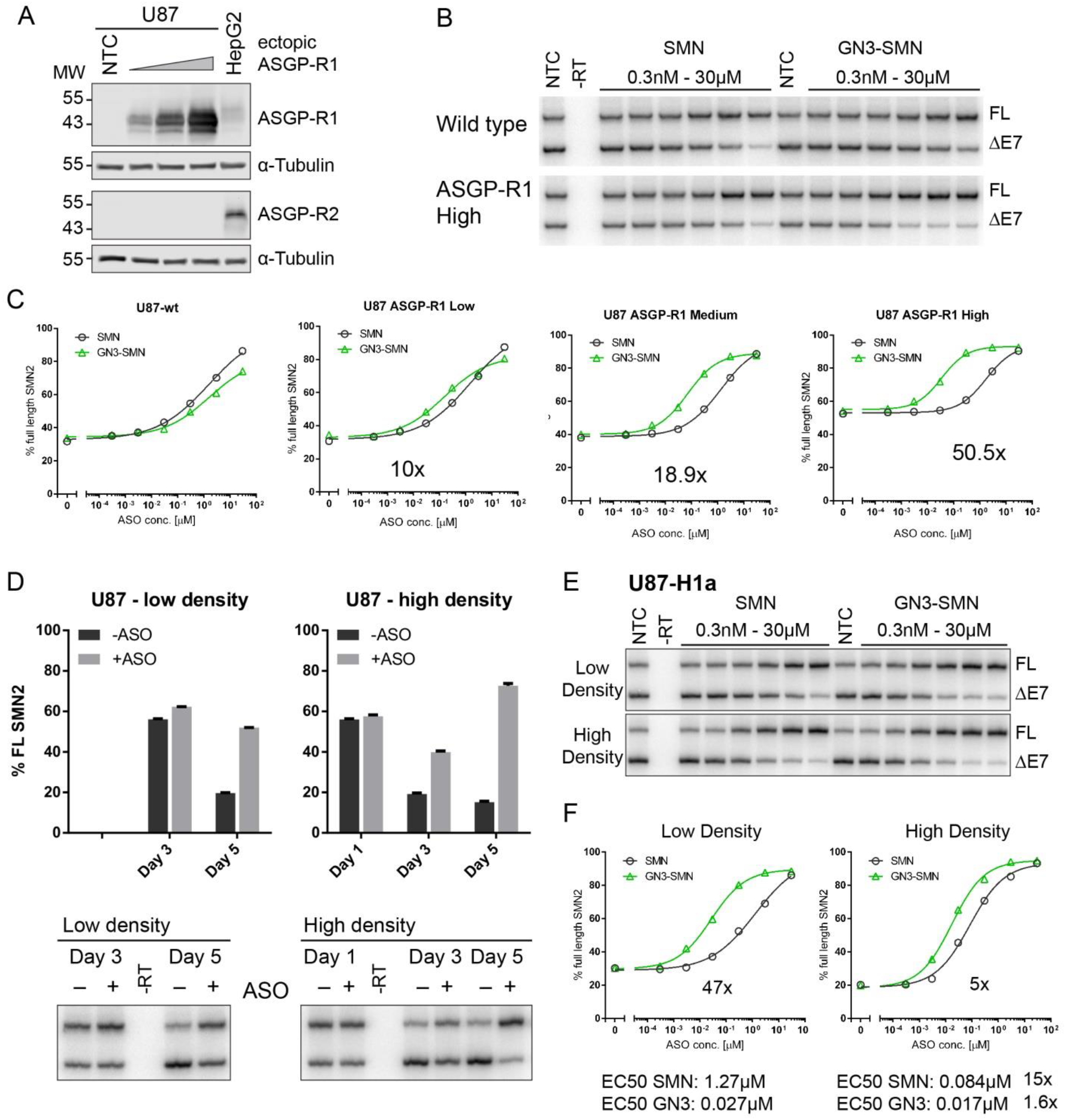
GN3-ASO activity positively correlates with ASGP-R1 expression level. (A) U87 cells were transduced with various titers of ASGP-R H1a retrovirus. Western blot shows increased expression of the ASGP-R1 isoform in U87 cells. (B) Representative radioactive gels of U87 and U87-H1a-high cells treated with SMN and GN3-SMN ASOs. Top band shows full-length *SMN2*, bottom band shows *SMN2* Δ exon 7 (DdeI digested). (C) Dose-response curves of ASO-treated U87 cells expressing increasing levels of ASGP-R1. (D) *SMN2* exon 7 inclusion in U87 cells is time- and cell-density-dependent. The proportion of full-length *SMN2* transcript decreases with time and cell density. Exon 7 inclusion of U87 cells treated with 300 nM SMN-ASO increases over time and in high-density conditions. Representative radioactive gels are shown below the bar chart. N=3 independent experiments; bars represent mean ± SE. (E) Radioactive gels showing *SMN2* splicing in low- and high-density U87-H1a cells treated with 0-30 μM SMN and GN3-SMN-ASO for 5 d. (F) Quantification of gels shown in panel E. Calculated EC50 doses are indicated below the graphs.

In the dose-response experiments described above, we noticed an increase in the baseline expression of FL-SMN2 in U87-high cells, compared to U87-wt and U87-low cells, which may influence ASO potency (Fig. 2C). *SMN2* splicing was shown to be influenced by the pH of the culture medium.^26^ U87-high cells were less confluent at the time of the analysis, and the medium pH was higher, likely because of the lower growth rate, compared to wild-type cells. We therefore looked at the dynamics of *SMN2* exon 7 splicing in a time- and cell-density-dependent manner, and how this dynamic influences ASO potency. As expected, exon 7 inclusion decreased markedly over time in untreated U87-wt cells, and this reduction took place earlier when the cells were seeded at a 4-fold higher density (Fig. 2D, -ASO). U87-wt cells treated with 300 nM SMN-MOE ASO by free-uptake showed increased exon 7 inclusion, compared to the untreated control, at all time points (Fig. 2D, +ASO). Furthermore, the effect of the ASO increased with time, likely due to slow and continuous cellular uptake and endosomal release. However, the effect of the ASO was also greater in cells seeded at a higher density, even at the same time point (compare Fig. 2D, day 5). This result is intriguing, because in low-density cultures there are fewer cells competing for the ASO, and the FL-SMN2 baseline is higher, which should favor ASO-induced exon 7 inclusion. The observed effect must therefore be caused by intrinsic changes in U87 cells cultured at a higher density, resulting in increased internalization or endosomal release. Interestingly, we did not observe a cell-density-dependent effect on the potency of unconjugated SMN-ASO in HEK293T cells (Fig. S2) or A172 cells (data not shown).

Next, we tested whether GN3-mediated delivery of ASO is also influenced by cell density. We integrated ASGP-R1 into the genome of U87 cells by lentiviral transduction to generate a stable cell line, and exposed low- and high-density cultures to unconjugated and GN3-conjugated ASOs at a final concentration of 0-30 μM for 5 days. Interestingly, the potency of unconjugated ASOs was 15-fold greater in U87-H1a cells grown at high density (Fig. 2E). The potency of GN3-conjugated ASOs, in contrast, was similar in low- and high-density cultures, with an EC50 of 27 nM and 17 nM, respectively (1.6-fold difference).

### Ectopic expression of ASGP-R in HepG2 cells increases GN3-ASO activity in 3D culture

ASGP-Rs are naturally expressed in hepatocytes, and promote GN3-mediated ASO-uptake *in vivo*. In hepatocellular carcinoma (HCC) cell lines, however, ASGP-R-mediated ASO delivery was reported to be inefficient, which may be explained by lower receptor levels in HCC cells.^23^ When we tested *SMN2* exon7 inclusion in HepG2 cells exposed to ASOs in the culture medium for 5 days, we found that unconjugated and GN3-conjugated ASOs performed very similarly (Fig. 3A). To show whether increased receptor expression improves GN3-ASO activity, we established HepG2 cells with stable expression of ASGP-R1 and R2, alone and in combination (Fig. 3B). Interestingly, overexpression of ASGP-receptors did not enhance GN3-ASO activity in HepG2 cells (Fig. 2C). GN3-ASOs did result in a slight increase of FL-SMN2, compared to unconjugated ASOs, in both wild-type and ASGP-R-expressing cells (46.9±5.5% and 53.8±6.7% FL-SMN2 in HepG2-wt, respectively, *P*=0.042). However, overexpression of receptors alone or in combination did not enhance GN3-ASO activity in HepG2 cells (Fig. 3C). Immunostaining of HepG2 cells for presence of the ASO in cells exposed to 300 nM ASO for 24 hrs is consistent with this finding: GN3-ASO staining was qualitatively indistinguishable between HepG2-wt and HepG2-H1a cells (Fig. 3D), which is drastically different from the results obtained in U87 cells (compare to Fig. 1G).

**Figure 3.**
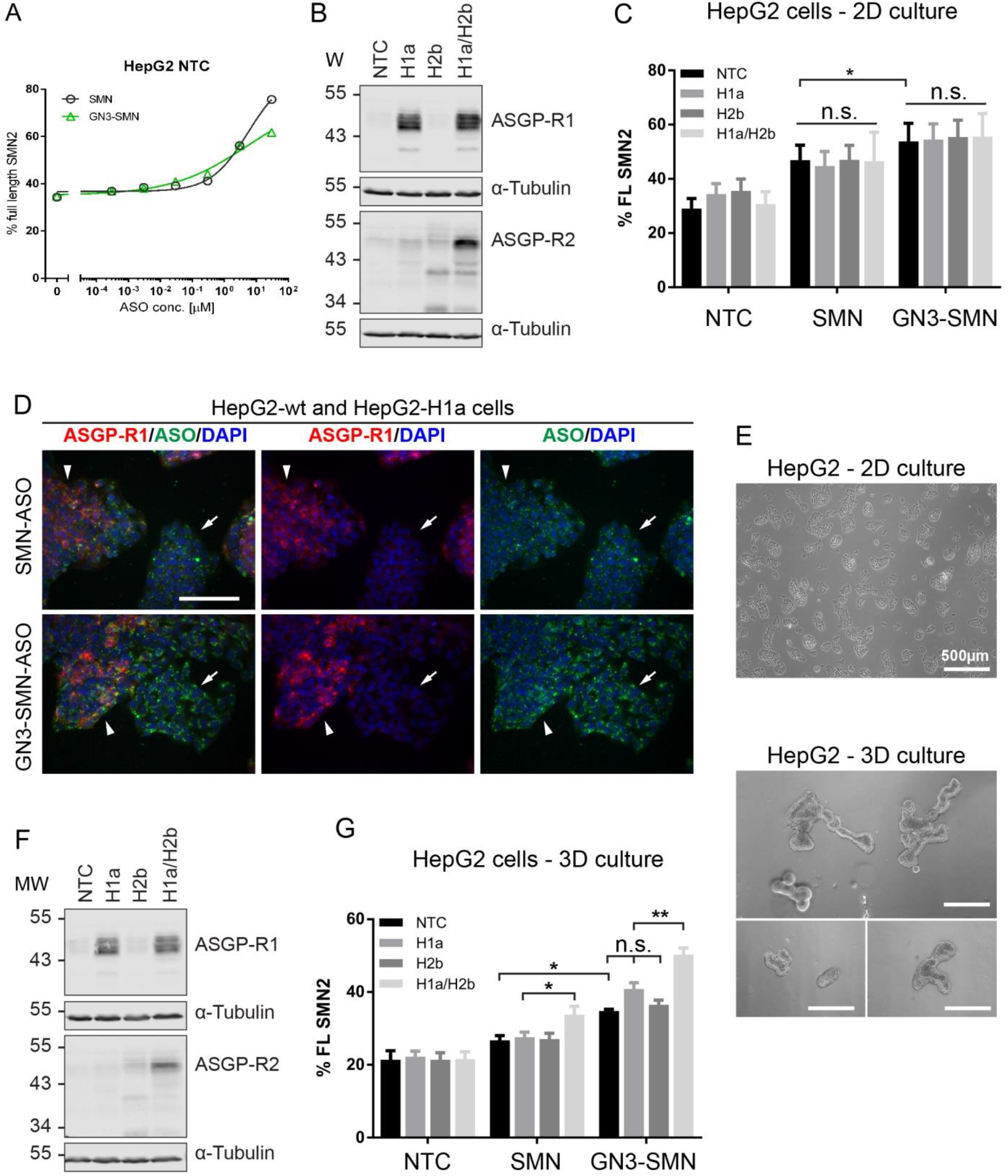
Ectopic expression of ASGP-R in HepG2 cells increases GN3-ASO activity in 3D culture. (A) Quantification of full-length *SMN2* mRNA in HepG2 cells treated with 0-30 μM SMN and GN3-SMN-ASO for 5 d. (B) Western blots of stable 2D HepG2 cells expressing ASGP-R H1a and H2b alone or in combination. Tubulin served as a loading control. (C) Quantification of full-length *SMN2* mRNA in 2D-HepG2 cells treated with 300 nM SMN and GN3-SMN-ASOs for 5 d. Ectopic expression of ASGP-R1 and/or 2 had no effect on ASO efficacy in 2D HepG2 cultures. (D) HepG2-wt (arrow) and HepG2-H1a cells (arrowhead) were cultured together and exposed to 300 nM SMN and GN3-SMN ASO for 24 hrs. Immunofluorescence staining shows ASGP-R1 (red), ASO (green) and nuclei (blue). HepG2-wt cells are ASGP-R1-positive, but the staining is very weak. Scale bar, 100 μm. (E) Phase-contrast micrographs of HepG2 cells grown attached on standard tissue-culture plates (top) or in 3D cultures on ultralow-attachment plates (bottom). Scale bars, 500 μM. (F) Western blots of stable 3D HepG2 cells expressing ASGP-R H1a and H2b alone or in combination. Tubulin served as a loading control. (G) Quantification of full-length *SMN2* mRNA in 3D-HepG2 cells treated with 300 nM SMN and GN3-SMN-ASOs for 5 d. Exon 7 inclusion upon GN3-SMN-ASO treatment is significantly greater in 3D HepG2 cells overexpressing ASP-R. N=3 independent experiments, bar graphs represent mean +/- S.E.; * *P* < 0.05; ** *P* < 0.01; NTC, no-treatment control; n.s., not significant.

HepG2 cells cultured in ultralow-attachment plates aggregate to form three-dimensional clusters of cells (Fig. 3E), and maintain ASGP-R expression (Fig. 3F). We delivered ASOs to these cultures at a final concentration of 300 nM by free-uptake for 5 days, before analyzing *SMN2* exon7 inclusion (Fig. 3G). In this case, HepG2-H1a cells treated with GN3-ASO showed slightly higher FL-SMN2 expression (40.8% FL-SMN2) compared to HepG2-wt (34.8% FL-SMN2) and HepG2-H2b cells (36.5% FL-SMN2). This increase between HepG2-H1a and HepG2-wt or HepG2-H2b was not statistically significant. However, in cells expressing both ASGP-R isoforms (H1a and H2b) in combination, GN3-ASOs performed significantly better than in cells expressing ASGP-R1 alone (50.3± 1.8% and 40.8±1.7% FL-SMN2 respectively, P=0.0023). Interestingly, ASGP-R H1a and H2b expressed together in HepG2 cells also increased the effect of unconjugated SMN-ASOs (Fig. 3G). This type of effect was reported before and is specific for PS-modified ASOs,^23, 24^ yet we did not observe it in U87 or HT1080 cells expressing the ASGP-Rs (Fig. 1F and S3).

### ASGP-R1 glycosylation state does not affect GN3-ASO mediated uptake

Next, we investigated the effects of posttranslational modifications on ASGP-R activity. ASGP-R1 protein migrates as several discrete bands by SDS-PAGE, which are largely non-overlapping when comparing U87 and HepG2 cells (Fig. 4A). When we expressed human ASGP-R1 in additional cell lines (Huh7, A172, LN308, HT1080 and 293T cells) we observed a wide variety of bands, even in cells of similar origin (compare HepG2 and Huh7 hepatic cells, and U87 and A172 glioblastoma cells) (Fig. 4B). This could potentially explain why ASGP-R1 performs better in some cell lines than others. Human ASGP-R1 has two Asn-residues that are glycosylated, N79 and N147.^27^ We hypothesized that the different bands might represent ASGP-R protein at various glycosylation stages, either because of incomplete processing (localized in the ER) or because some cell types do not express the required glycosyltransferases. To test whether proteins are alternatively glycosylated, we digested protein samples with Peptide-N-Glycosidase F (PNGase F), an endoglycosidase that catalyzes the deglycosylation of most N-linked glycoproteins.^28^ ASGP-R1 protein in untreated whole-cell lysates of U87-H1a and HepG2-H1a cells had different electrophoretic mobilites (Fig. 4C). Upon PNGase treatment, all slower-migrating bands disappeared and coalesced into a single, faster-migrating band. This result confirms that there are multiple glycosylation states of each protein, as well as cell-type-specific glycosylation patterns.

**Figure 4.**
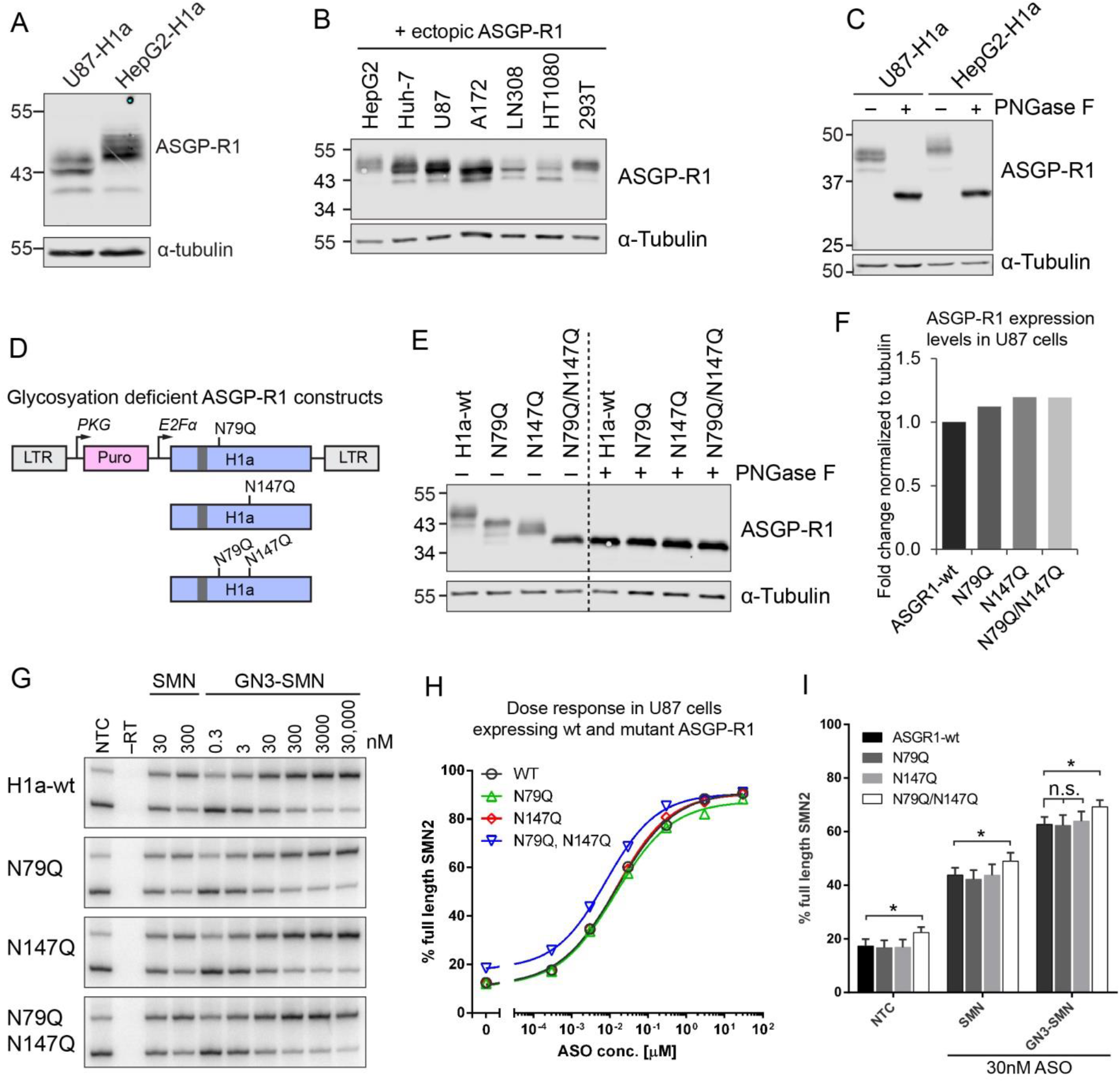
ASGP-R1 glycosylation state does not affect GN3-ASO efficacy. (A) Western blot of U87 and HepG2 cells overexpressing ASGPR-H1a. Multiple bands with different electrophoretic mobilities are observed for both cell lines, likely due to variable posttranslational modifications. (B) Ectopic expression of ASGP-R1 in other cell lines also shows that ASGP-Rs are differentially modified in each cell line. (C) U87-H1a and HepG2-H1a cell-lysates were deglycosylated with Peptide-N-Glycosidase F (PNGase F). ASGP-R1 protein in untreated whole-cell lysates of U87-H1a and HepG2-H1a cells migrates at different apparent molecular weights. Upon PNGase treatment, all slower-migrating bands disappear and coalesce into a single, faster-migrating band, suggesting multiple glycosylation states of each protein, as well as cell-type-specific glycosylation patterns. (D) Glycosylation-deficient ASGP-R1 cDNA in lentiviral expression vectors. Glycosylated asparagines N79 and N147 were mutated to glutamines, either alone or in combination. (E) U87 cells stably expressing glycosylation-deficient ASGP-R1 mutants. Bands of Asn-mutant ASGP-R1 migrate faster, consistent with deficient glycosylation. PNGase-F-digested protein bands (deglycosylated) co-migrate, similar to the double mutant. (F) Quantification of protein bands in panel E. Bars show fold-change compared to ASGP-R1 wild-type-expressing U87 cells, normalized to tubulin. (G) Radioactive RT-PCR showing *SMN2* splicing in stable U87 cells expressing wild-type and Asn-mutant ASGP-R1, treated with 0-30 μM SMN and GN3-SMN-ASO for 5 d. (H) Quantification and dose-response curves from data in panel G. (I) Cells were treated with 30 nM SMN-ASO and GN3-SMN ASO for 5 d, in triplicates for statistical analysis. Bar graphs represent mean +/- S.E.; * *P* < 0.05; ** *P* < 0.01; NTC, no-treatment control; n.s., not significant.

To test whether the glycosylation state affects ASGP-R activity, we mutated both Asn-residues to Gln, either alone or in combination, and cloned these glycosylation-deficient mutants into a lentiviral expression vector to generate stably transduced U87 cell lines (Fig 4D-E). Western blotting of U87 cells expressing the Asn-mutants shows multiple bands. Upon deglycosylation with PNGaseF, all protein species co-migrated as a single band (Fig 4E). Furthermore, the expression level of the Asn-mutant receptors was similar in the different stable cell lines (Fig. 4F). When we tested the effect of GN3-conjugated ASOs in U87 cells expressing Asn-mutant receptor isoforms, we found that the glycosylation state of the receptor had very little to no influence on ASO uptake and activity (Fig. 4G-H). Mutant receptors performed as well as fully glycosylated, wild-type ASGP-R1 (Fig. 4I). The improved potency of the N79Q/N147Q double mutant (1.9 fold) was statistically significant; however, the baseline expression of FL-SMN2 was also increased in double-mutant cells, which could potentially contribute to this effect.

### GN3-conjugated ASOs can be delivered to ASGP-R1-expressing tumor xenografts *in vivo*

Tumor xenografts are notoriously difficult to target by antisense therapy.^29^ A primary goal of this study, therefore, was to bypass the natural uptake route by ectopically expressing ASGP-R1, and determine whether ASGP-R1-expressing tumor cells can be specifically and efficiently targeted by GN3-ASOs *in vivo*. We used a subcutaneous U87 xenograft model to test receptor-specific targeting of GN3-ASOs. We subcutaneously implanted human U87 cells expressing isoform H1a, and control cells expressing isoform H2b, into immunocompromised NSG mice, and administered a total of four subcutaneous ASO doses at 200 mg/kg/week, starting 10 days post-implantation (Fig 5A). After a two-week treatment regimen, we harvested tumor, liver, and kidney tissues to visualize the ASO distribution by IHC, and to measure *SMN2* exon 7 inclusion by RT-PCR (Fig. 5B-E). Saline and an unconjugated random-sequence control ASO served as negative controls. We stained all the samples with H&E, which did not reveal obvious abnormalities or signs of toxicity (Fig. S4). U87-H1a and U87-H2b tumor samples processed for IHC-staining confirmed the expression of these two isoforms, respectively (Fig. 5B). Mouse liver samples served as a control, and naturally express both isoforms.

**Figure 5.**
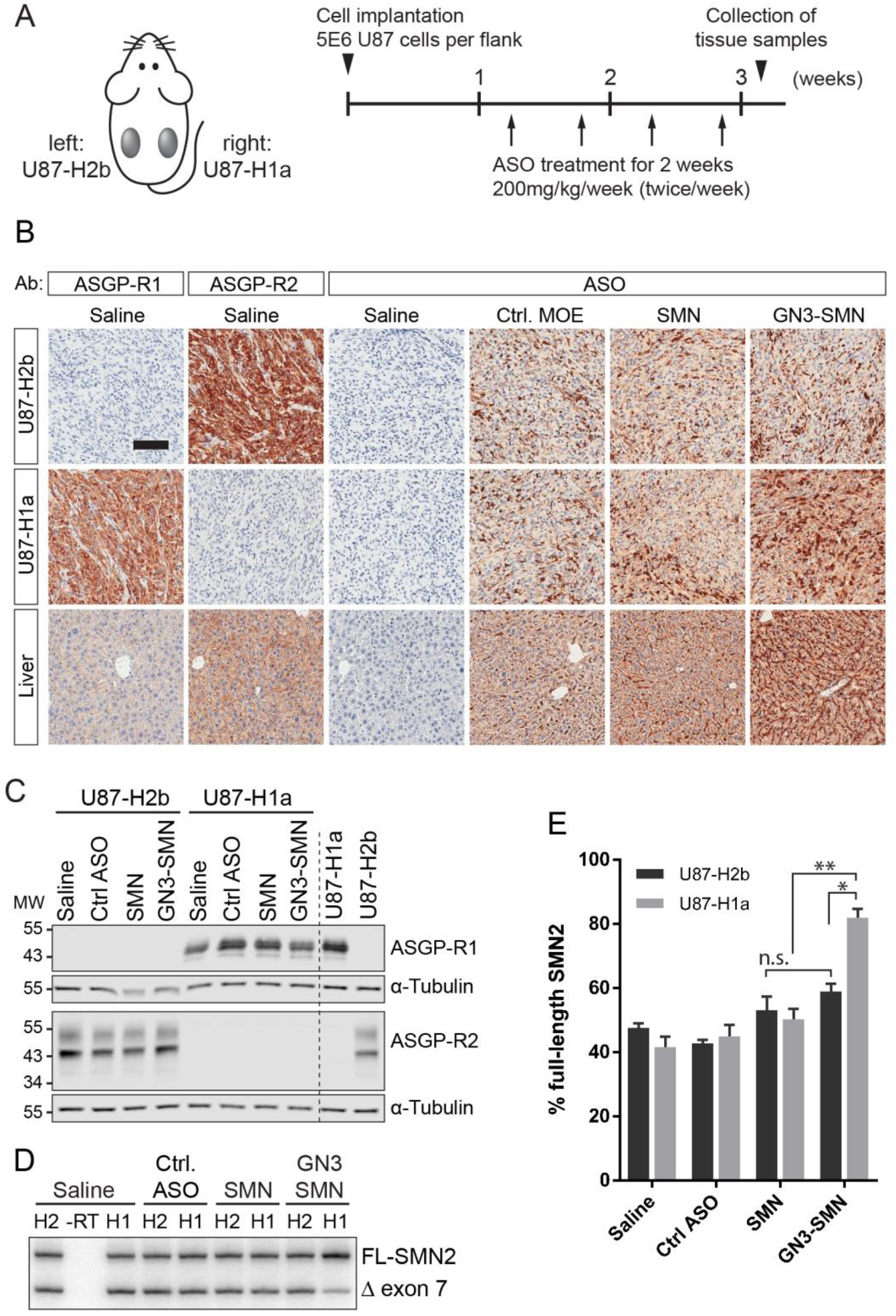
Delivery of GalNAc-conjugated ASOs to tumor xenografts. (A) Schematic of the tumor xenograft model and ASO dosing schedule of tumor-bearing NSG mice. 5×10^6^ U87-H2b cells (control) and U87-H1a cells (active receptor) were subcutaneously implanted into the left and right flank of adult NSG mice, respectively. Three animals per group were treated with saline, control ASO, SMN-ASO, and GN3-SMN-ASO at 200 mg/kg/week, with two s.q. injections per week. Three days after the last injection, tumor, liver and kidney samples were collected and processed for RNA, protein and histological analysis. (B) Representative pictures of IHC analysis of ASGP-R1/2 expression and ASO localization in U87-H2b, U87-H1a, and liver sections, after 2 weeks of ASO treatment. The rabbit-anti-ASO antibody (Ionis Pharmaceuticals) recognizes the ASO’s phosphorothioate backbone. Scale bar, 100 μm. (C) Western blot analysis of U87-H1a and U87-H2b tumors confirms ASGP-R1 and 2 expression. Cultured U87 cells expressing ASGP-R1 and 2 express comparable receptor levels. (D) *SMN2* exon 7 inclusion ratio in saline- and ASO-treated U87-H1 and U87-H2 control tumors was quantified by radioactive RT-PCR. (E) Quantification of results in panel C. N=3 animals/treatment; bar graphs represent mean ± SE; * *P* < 0.05; ** *P* < 0.01; n.s., not significant.

Sections stained for ASO localization with a phosphorothioate-backbone-specific antibody showed a signal in all ASO-treated samples, but not in the saline-treated controls. Unconjugated control and SMN ASOs showed similar signal intensities in U87-H1a and U87-H2b tumors. GN3-conjugated SMN-ASOs, however, showed much stronger staining in U87-H1a cells than in U87-H2b cells. Furthermore, the signal was also stronger than with unconjugated ASOs, both in U87-H1a tumors and mouse liver, but not in U87-H2b tumors. In mouse kidney, both unconjugated and GN3-conjugated ASOs were internalized by proximal convoluted tubules (Fig. S4). We used Western blotting to measure ASGP-R expression in U87 tumors, which was comparable to that in cultured cells (Fig. 5C). The U87-H1a-specific targeting of GN3-ASOs also correlated with significantly higher *SMN2* exon 7 inclusion in U87-H1a tumors than in U87-H2b tumors (Fig. 5D-E). We also tested a different dosing schedule (more frequent injections and 20% more ASO), which slightly increased FL-SMN2 levels in U87-H1a tumors. In conclusion, GN3-mediated delivery can be used to test therapeutic splice-switching ASOs against targets of interest in non-hepatic cells *in vivo*, by overexpressing ASGP-R1, which is advantageous in the case of difficult-to-target disease models, such as cancer.

## Discussion

RNA-targeting antisense drugs have great clinical potential; however, efficient delivery of ASO to the intended target tissue remains a challenge in many cases. There is a concerted effort in the field to find new ASO formulations, such as lipid nanoparticles or receptor/ligand pairs, which will allow efficient and tissue-specific targeting. The advantages of such an approach are threefold: (1) tissue-specific targeting increases the potency of the ASO by increasing its availability where it engages its therapeutic target; (2) potential sequence-specific or chemistry-class off-target effects would be limited to the targeted tissue; and (3) efficient internalization in the target tissue, together with long ASO half-life, means that lower and less frequent dosing would be required, further improving drug safety.

One receptor/ligand system whose clinical importance has already been demonstrated is ASGP-R. It is expressed in hepatocytes and forms multimeric units to bind and internalize GN3-conjugated ASOs. Out of all five membrane-bound isoforms, ASGP-R H1a has been shown to be required to bind the GN3-ligand.^24^ However, until very recently, it was not known whether it acts cooperatively with ASGP-R2 isoforms.^23^ Here we confirm that expression of isoform H1a alone is sufficient to enhance the activity of GN3-conjugated ASOs in multiple cancer cell lines. However, the effects varied markedly in different cell lines and under different culture conditions. Given that ASGP-R1 protein migrates as multiple apparent-molecular-weight bands that vary across cell lines, we considered that alternative posttranslational modifications may affect receptor activity. Asialoglycoprotein receptors are glycosylated, phosphorylated, and fatty-acylated, and receptors on the cell surface are mostly hypophosphorylated and hyperglycosylated.^30^ However, we found that mutating ASGP-R1 glycosylation sites had little to no effect on its activity in U87 cells. We cannot rule out an effect of the phosphorylation state of the receptor, which we did not test in this study. ASGP-R1 protein species appear to be identical in 2D and 3D cultures of HepG2 cells. However, the observed difference in uptake between unconjugated and GN3-conjugated ASOs is greater in 3D culture, which may be due to external factors, such as altered cell polarity, affecting ASO internalization or intracellular trafficking.

In addition to enhancing GN3-ASO uptake, ASGP-R has also been reported to increase the effect of unconjugated P=S-modified ASOs *in vitro* and *in vivo*, albeit to a much lower extent than with GN3-ASOs.^23^ In contrast to these studies, our results did not reveal a measurable impact of ASGP-R1 expression on SMN-ASO activity in A172, HEK293T, and HT1080 cells. However, we did see a small but significant impact in HepG2 cells, but only when both ASGP-R H1a and H2b were expressed together, and only in 3D culture. ASOs used in previous reports were gapmers with several unmodified DNA bases, whereas the SMN-ASOs used in the present study were uniformly MOE-modified, which could account for the discrepancy.

Another possible explanation for the above discrepancy is the duration of treatment. Cells treated with gapmer ASOs are frequently analyzed after a 24-hour exposure to the ASO, versus a 5-day exposure in the case of SMN splice-modulating ASOs. The different time courses reflect in part the destruction of RNA in the case of gapmers, versus the gradual change in the proportion of two isoforms, in the case of splice-modulating ASOs. GN3-ASOs are internalized rapidly, as shown here and by others. However, time-course experiments with SMN-ASOs show a gradual increase in *SMN2* exon 7 inclusion over a 5- day treatment period in different cell lines, likely due in part to slow endosomal release becoming a limiting factor in ASO delivery. Assessing splice-modulating ASO-activity over time is difficult, because endogenous *SMN2* splicing is very dynamic in cell culture, and is cell-type-specific. For example, in untreated U87 cells, FL-SMN2 decreases over time; in HEK293T cells, FL-SMN2 levels go up; and in A172 cells, *SMN2* splicing remains largely unchanged. *SMN2* splicing is influenced by altered expression of splicing factors, which is regulated by environmental factors, such as the pH of the medium.^26^

Perhaps the most unexpected result in this study is that the activity of unconjugated ASOs was heavily influenced by cell density in U87 cells. Unconjugated SMN-ASOs were 15-fold more potent in high-density cultures than in low-density cultures, after a 5-day treatment period. The ASO phosphorothioate backbone has been shown to bind proteins, which may facilitate ASO internalization via unknown receptors.^31^ Environmental changes in high-density U87 cultures, such as lower medium pH, faster growth-factor depletion, and increased cell-cell contacts, may result in expression or conformational changes of unidentified cell-surface receptors required for ASO uptake. Alternatively, environmental and/or physiological changes may also affect endosomal release or intracellular ASO trafficking. However, this latter explanation seems less likely, because ASO delivery via ASGP-R did not appear to be affected by cell density, showing more consistent dose-dependent *SMN2* exon 7 inclusion.

The growth conditions of cancer cells in tumor mouse models are very different from cell-culture conditions. In addition to cellular uptake itself, ASO delivery to cancer cells *in vivo* is also influenced by plasma concentration and tumor vascularization. Furthermore, wild-type and ASGP-R-expressing tumor cells compete with other cell types, including hepatocytes, which take up and clear ASOs rapidly. Successful delivery of unconjugated and GN3-conjugated ASOs to cancer cells was therefore not guaranteed. Consistent with this assumption, FL-SMN levels increased only marginally in unconjugated-ASO-treated tumors, even at doses far above those required to correct *SMN2* splicing in the CNS or in hepatocytes of SMN2-transgenic mice. GN3-conjugated ASOs, in contrast, were efficiently targeted to ASGP-R-expressing cancer cells, resulting in robust splice-switching *in vivo*.

Another potential application of the receptor/ligand-system could be to ectopically express ASGP-R in specific tissues by lentiviral transduction or transplantation of ASGP-R-expressing stem cells. For example, one could use receptor-expressing muscle satellite cells to target ASOs specifically to regenerating muscle, or express ASGP-R in a subpopulation of cells in the CNS with a viral vector, followed by ASO treatment. This would not only allow testing of therapeutic ASOs, but could also serve to generate a disease phenotype associated with a splicing defect in a specific cell population, expanding the molecular toolbox to study splicing in health and disease.^32^

In conclusion, our work shows that GN3-mediated targeting of ASOs, combined with ectopic ASGP-R expression, is an effective tool for proof-of-principle studies to test novel ASO-mediated splice-switching therapies, until more efficient and specific targeting methods become available.

## Material and Methods

### Animals and tumor model

Nod-Scid-gamma (NSG) immunocompromised mice (The Jackson Laboratory strain #005557) were housed in vented cages and bred in-house. U87 tumor xenografts were established by injecting 5×10^6^ U87 cells (50,000 cells/μL in HBSS) subcutaneously into the flanks of adult NSG mice. Once tumors were palpable (7-10 days post-transplantation), animals were treated with ASOs delivered by s.q. injection at 200-250 mg/kg/week, with 2-5 injections per week. All animal protocols were performed in accordance with Cold Spring Harbor Laboratory’s Institutional Animal Care and Use Committee (IACUC) guidelines.

### Plasmids

Full-length human *ASGR1* and *ASGR1* cDNA sequences were amplified from HepG2 cDNA samples and cloned into a pMSCV retroviral backbone^33^ by conventional cloning, and into a lentiviral backbone containing a puromycin selection marker (kindly provided by Scott Lyons, CSHL) using Gibson Assembly cloning (NEB, Ipswich, MA). Retro- and lentiviruses were produced in HEK-293T/17 cells by co-transfecting viral constructs with psPAX2 and VSVG. Viral supernatant was collected 48-72 hrs post-transfection, filtered and stored at -80 °C. To generate stable cell lines, U87 and HepG2 target cells were infected with lentiviral particles overnight, in the presence of 8 μg/mL polybrene (Sigma, St. Louis, MO) and selected using 1-2 μg/mL puromycin (Sigma, St. Louis, MO) for at least 2 weeks. Cell lines expressing both ASGP-R H1a and H2b isoforms were transduced individually on two consecutive days. Retroviral particles were only used for transient experiments.

### Cell culture

U87 MG glioblastoma cells, A172 glioblastoma cells, HT1080 fibrosarcoma cells, and HEK-293T/17 cells were maintained in DMEM (Corning, Manassas, VA), supplemented with 10% FBS (Seradigm VWR, Radnor, PA) and 1% penicillin/streptomycin. HepG2 hepatocellular carcinoma cells were maintained in EMEM (ATCC, Manassas, VA), supplemented with 10% FBS and 1% penicillin/streptomycin at 37 °C/5% CO2. 3D-cultures were induced by seeding cells in ultra-low attachment plates (Corning, Kennebunk, ME).

### Antisense oligonucleotides

All ASOs used were uniformly modified with MOE sugars, phosphorothioate backbone, and 5’-methyl cytosine, and are listed in Table 1. MOE synthesis, purification, and quantification were done as described.^34^ GalNAc-conjugated ASOs were synthezised, purified, and quantified as described.^18^ The GalNAc moiety was conjugated to the 5’-end of the ASO with a trishexylamino (THA)-C6 cluster. All ASOs were dissolved in water and stored at -20 °C. The sequences of all ASOs used in this study are provided in Supplementary Table 1. ASOs at concentrations ranging from 0.3 nM to 30 μM were delivered by free uptake for 1-5 days *in vitro*.

**Table 1.**
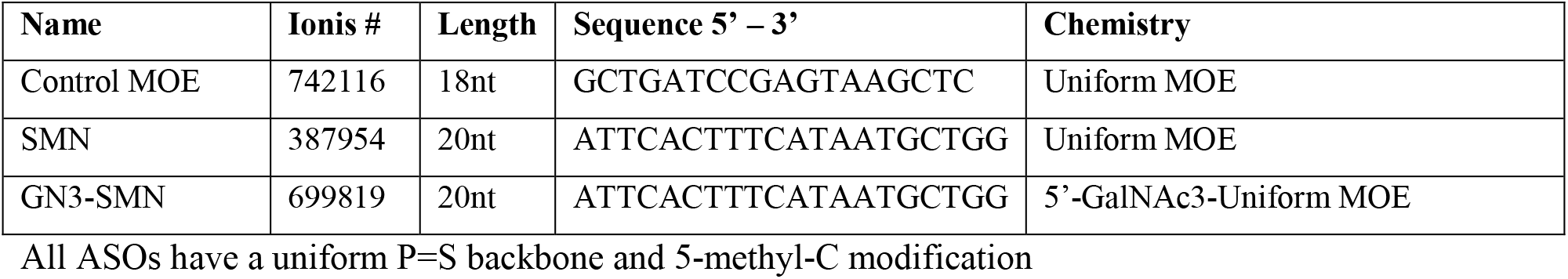
Antisense oligonucleotides used in this study.

### PCR

RNA was extracted from cells and tissue using Trizol reagent (Invitrogen, Carlsbad, CA), and reverse-transcribed using ImProm-II reverse transcriptase (Promega, Madison, WI). PCR was performed with AmpliTaq polymerase (Thermo Fisher, Foster City, CA) using human SMN-specific exon 6 Fwd 5’-ATAATTCCCCCACCACCTCCC-3’ and exon 8 Rev 5’-TTGCCACATACGCCTCACATAC-3’ primers at a final concentration of 250 nM. [α-32P]-dCTP radiolabeled PCR products were digested with DdeI (NEB, Ipswich, MA) for 2 hrs at 37 °C and separated on a 5% native polyacrylamide gel (Biorad, Hercules, CA), analyzed on a Typhoon 9410 phosphorimager (GE Healthcare) and quantified using Multi Gauge v2.3 (Fujifilm, Tokyo, Japan). The radioactive signal from each band was normalized to the G/C content to calculate relative changes in splice isoforms.

### Western Blotting

Cells and tissues were harvested, lysed in RIPA buffer supplemented with 1× complete protease inhibitor cocktail (Roche, Indianapolis, IN), and sonicated for 10 s (1s on/1s off). The protein concentration of the soluble fraction was determined by Bradford assay (Biorad, Hercules, CA) before adding Laemmli loading buffer. To assay the protein glycosylation state, the soluble fraction was denatured and treated with PNGase F (NEB P0809, Ipswich, MA) for 1 hr at 37 °C, according to the manufacturer’s instructions, before adding Laemmli loading buffer. Protein lysates were separated on denaturing polyacrylamide gels, transferred onto nitrocellulose membranes, and blocked with 5% milk in TBST. Membranes were incubated overnight at 4 °C with blocking buffer containing primary antibodies rb-ASGP-R1 (1:6000; ProteinTech 11739-1-AP), rb-ASGP-R2 (1:3000; Abcam ab200196, Cambridge, UK), ms-α-tubulin (1:10,000; Sigma T9026). After incubating the membranes with goat-anti-mouse and goat-anti-rabbit Licor IRDye 800 (green) and 680 (red) secondary antibodies (1:10,000; Licor, Lincoln, NE) in blocking buffer for 1 hr at room temperature, protein bands were visualized on an Odyssey imaging system (Licor, Lincoln, NE) and analyzed using ImageStudio and ImageJ. EZ-Run prestained markers (Fisher Scientific, Hampton, NH) or Precision Plus Protein Dual Color Standards (Bio-Rad, Hercules, CA) served as molecular-weight markers.

### Immunofluorescence

Cells were seeded onto 8-well culture slides (Falcon, Big Flats, NY), fixed with 4% PFA/PBS and permeabilized with 0.5% TritonX-100/PBS. Samples were blocked with 10% normal goat serum (NGS, Invitrogen, Rockford, IL) and 2% BSA in TBST for 1 hr at room temperature, and incubated with primary antibodies ms-ASGP-R1 (1:1000, Novus Biologicals MAB4394), rb-ASGP-R2 (1:500, Abcam ab200196) and rb-ASO (1:1000, Ionis Pharmaceuticals) in TBST supplemented with 2% NGS and 2% BSA at 4 °C overnight. Secondary antibodies used were AlexaFluor 488 (green) and 594 (red) (1:500, Invitrogen, Eugene, OR), and nuclei were stained with DAPI. Pictures were captured on a Zeiss Observer microscope. All images within the same figure panel were taken with the same exposure settings, and identically processed using Zeiss Zen software and Adobe Photoshop.

### Immunohistochemistry

Tissue samples were fixed in 4% PFA/PBS (EMS, Hatfield, PA) and paraffin embedded. 6-μm sections were deparaffinized and rehydrated. Endogenous peroxidase was blocked in H2O2/methanol for 15 min. Slides were then boiled in 10 mM sodium citrate/Tween for 5 min under pressure (for protein antigens), or treated with Proteinase K (Dako, Carpinteria, CA) for 8 min to unmask epitopes (for ASO staining). Slides were blocked for 30 min with peptide blocking solution (Innovex Biosciences, Richmond, CA) and incubated with the following primary antibodies for 1 hr at RT: rb-ASGP-R1 (1:500, ProteinTech 11739-1-AP), rb-ASGP-R2 (1:600, Abcam ab200196, Cambridge, UK), rb-ASO (1:10,000; Ionis Pharmaceuticals). The signal was visualized with HRP-labelled anti-rabbit polymer (DAKO, Carpinteria, CA) and DAB (DAKO, Carpinteria, CA). Slides were counterstained with Hematoxylin (Sigma, St. Louis, MO), mounted with Limonene mounting medium (Abcam, Cambridge, UK).

### Statistics and graph representation

Paired Student’s t-test was used to test statistical difference between untreated and ASO-treated samples, and between unconjugated and GN3-conjugated ASOs, *in vitro* and *in vivo*. EC50 values were determined by fitting a sigmoidal dose-response curve (variable slope) to the data, with bottom and top limits set to ≥0% and ≤100% full-length *SMN2*, respectively. Graphs were generated using Graphpad Prism and Microsoft Excel. Figures were compiled using Adobe Photoshop and Adobe Illustrator.

## Acknowledgements

We thank Scott Lyon (CSHL) for providing the lentiviral vector backbone, Michael Wigler (CSHL) for providing HT-1080 cells. We acknowledge support from NCI Program Project Grant CA13106 and the Histology Core CSHL Cancer Center Support Grant (CA045508).

## Author Contributions

Conceptualization, J.S. and A.R.K.; Investigation, J.S. and S.Q.; Writing – Original Draft J.S.; Writing – Review & Editing, J.S. A.R.K., F.R., C.F.B.; Resources, A.R.K., F.R., C.F.B.; Supervision, A.R.K.

---

**Figure S1.**
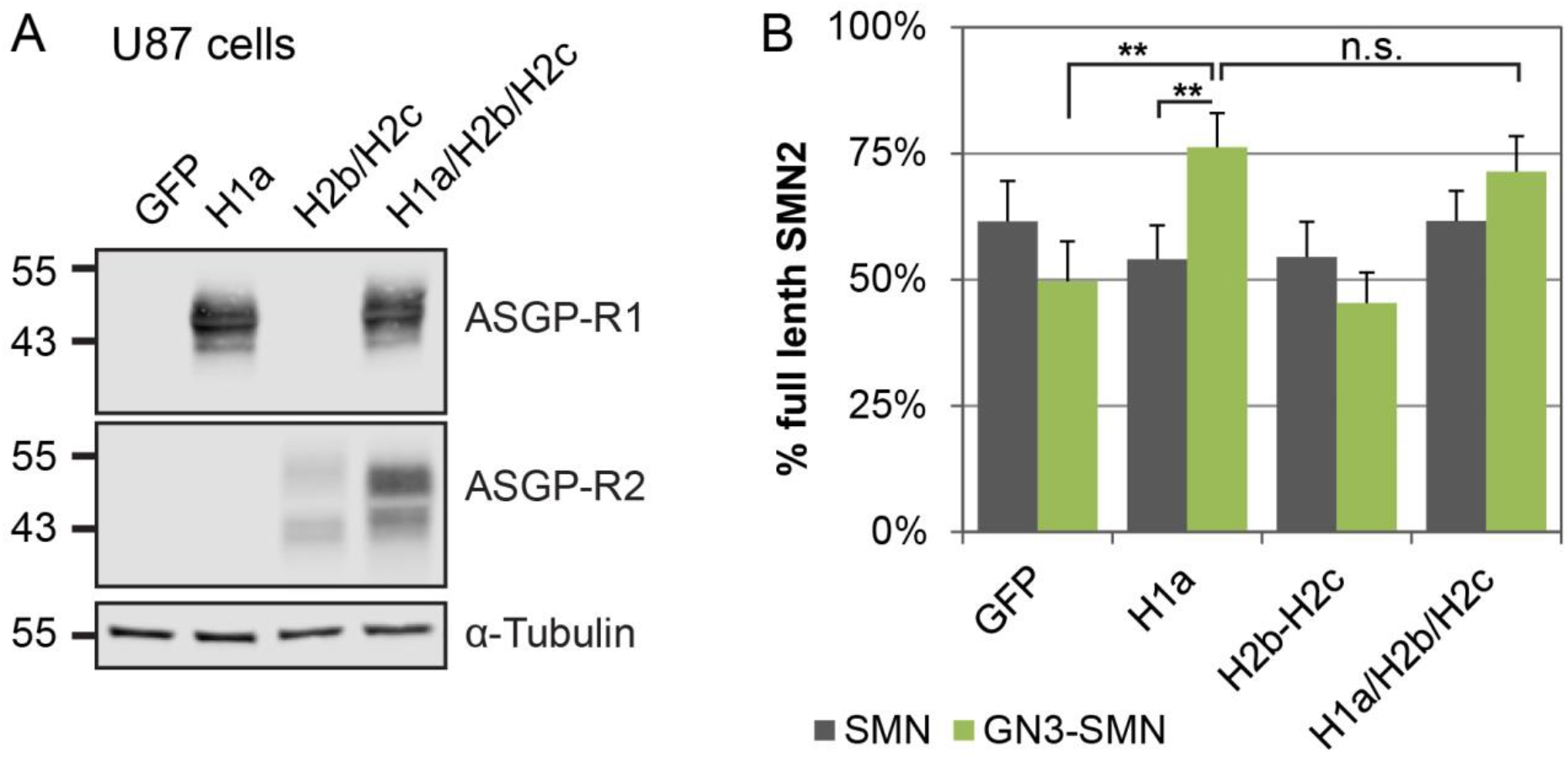
Additional ASGP-R isoform combinations tested in U87 cells. Expressing either isoforms H2b and H2c, or all three isoforms in combination, does not increase ASO activity in U87 cells, compared to cells expressing ASGP-R H1a alone. (A) Western blot confirming ectopic expression of ASGP-R isoforms in U87 glioblastoma cells. (B) Quantification of full-length *SMN2* mRNA in U87 cells treated with 300 nM SMN and GN3-SMN ASOs for 5 d by free-uptake. Bar graph represents mean ± SE of three independent experiments; ** *P* < 0.01; n.s., not significant.

**Figure S2.**
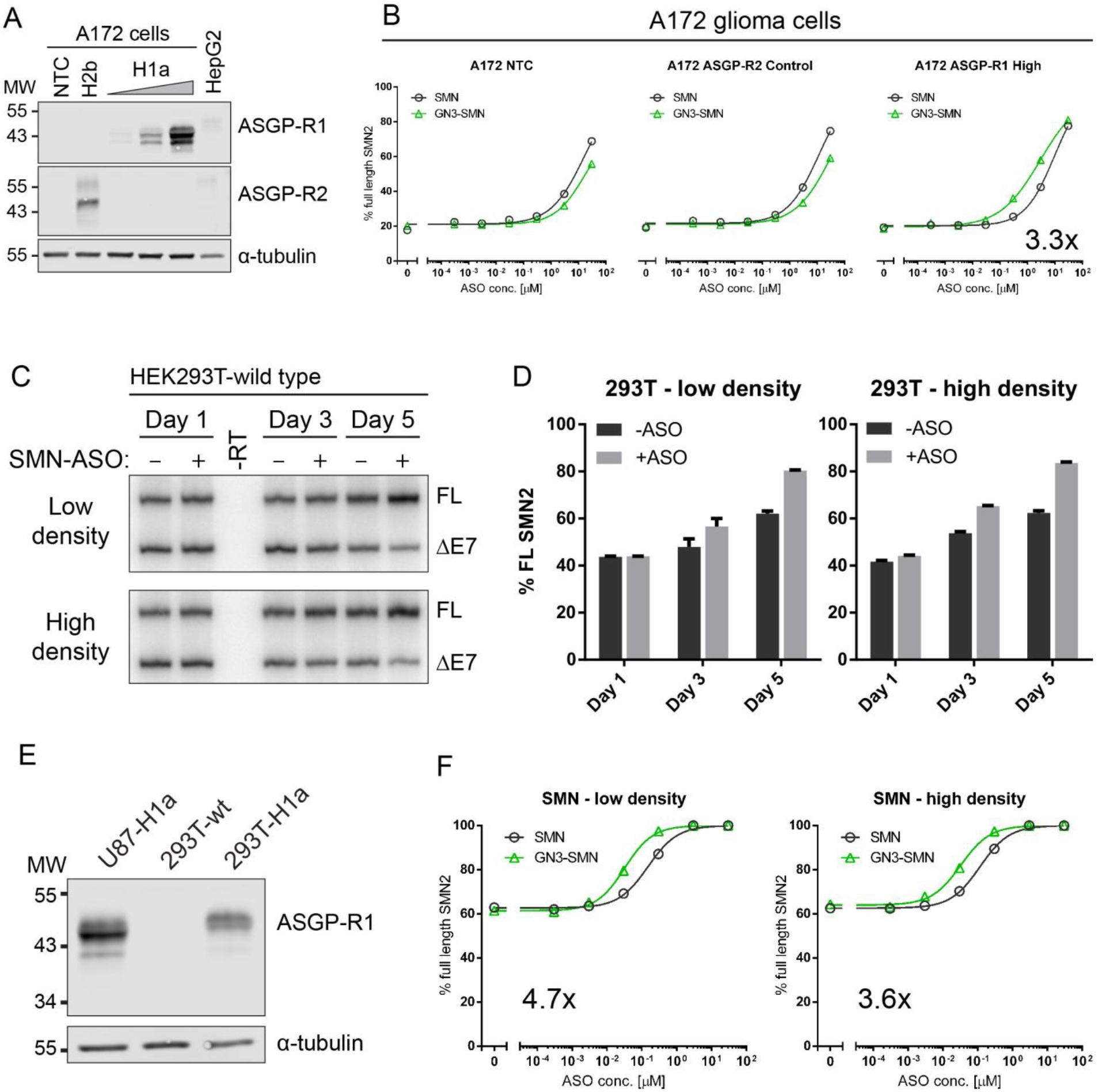
ASGP-R receptor activity in A172 and HEK293T cells. (A) Western blot of A172 cells expressing ASGP-R H2b, or increasing levels of ASGP-R H1a. HepG2 cells express both ASGP-R1 and 2 endogenously, and were included as a control. Tubulin served as loading control. (B) A172 cells treated with 0-30 μM SMN and GN3-SMN ASOs for 5 d by free uptake. Seven-point dose-response graphs represent full-length *SMN2* transcript that increases with ASO concentration. (C) Representative gel radiographs of low- and high-density HEK293T-wt cells treated with SMN-ASO for 1-5 d by free uptake. (D) Quantification of data in panel C. Bar graph represents mean ± SE of three independent experiments. (E) Western blot of HEK293T cells stably expressing ASGP-R1. Stable U87-H1a cells are shown for reference. (F) HEK293T-H1a cells treated with 0-30 μM SMN and GN3-SMN ASOs for 5 d by free uptake.

**Figure S3.**
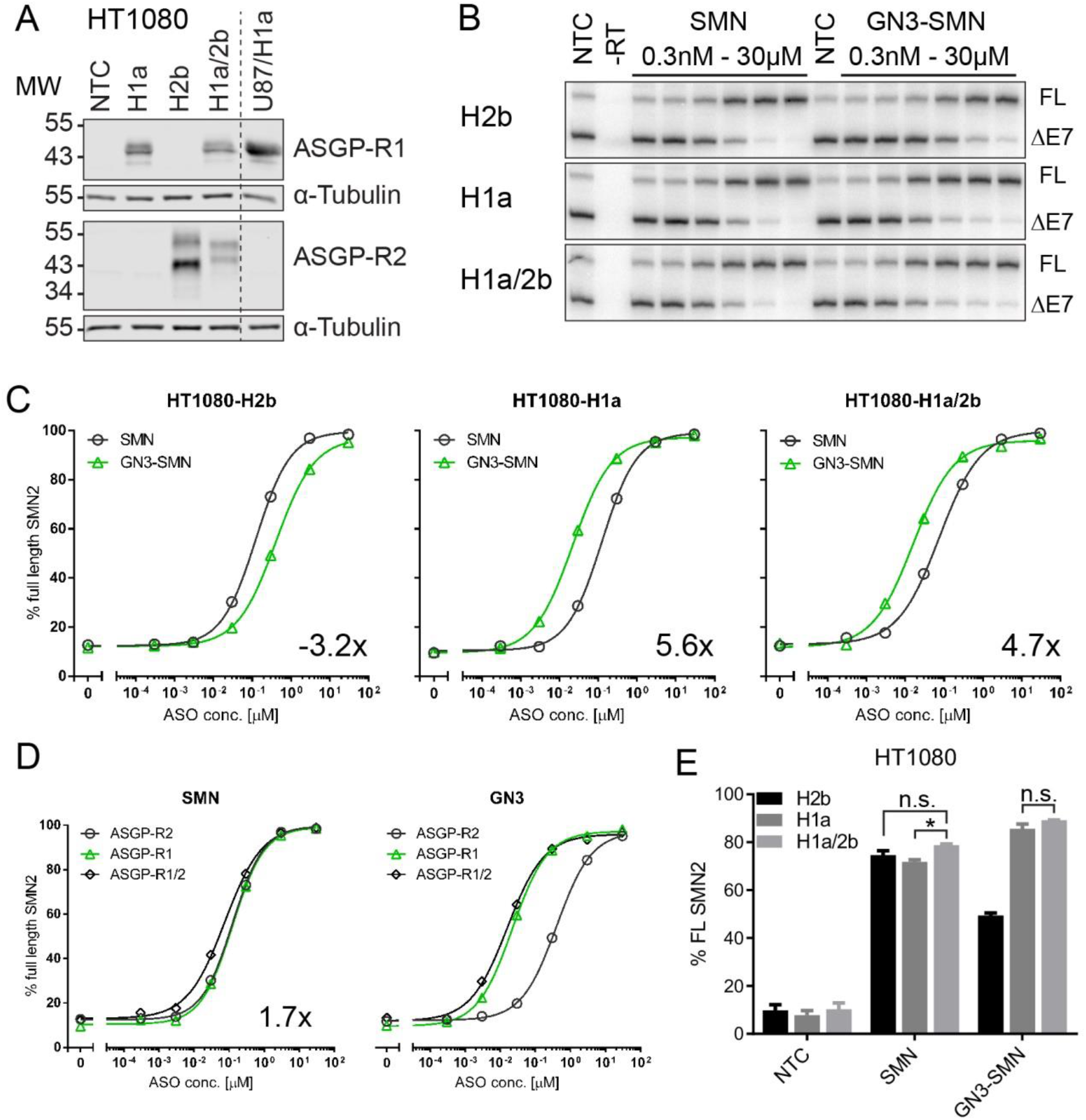
ASGP-R receptor activity in HT1080 fibrosarcoma cells. (A) Western blot of HT1080 cells expressing ASGP-R H1a and H2b alone and in combination. U87-H1a cells were included as a control, and tubulin served as loading control. (B) Representative radioactive gels of HT1080-H2b, -H1a, and -H1a/H2b cells treated with SMN and GN3-SMN ASOs. Top band shows full-length *SMN2*, bottom band shows *SMN2* Δ exon 7 (DdeI digested). (C) HT1080 cells treated with 0-30 μM SMN and GN3-SMN ASOs for 5 d by free uptake. Seven-point dose-response graphs represent full-length *SMN2* transcript that increases with ASO concentration. The fold-difference in EC50 is indicated in each panel. (D) Comparison of dose-response curves of SMN and GN3-SMN ASOs (E) Quantification of full-length *SMN2* mRNA in HT1080 cells treated with 300 nM SMN and GN3-SMN-ASOs for 5 d. Exon 7 inclusion upon GN3-SMN-ASO treatment was significantly greater in HT1080-H1a and HT1080-H1a/H2b cells. N=3 independent experiments, bar graphs represents mean +/- S.E.; * *P* < 0.05; NTC, no-treatment control; n.s., not significant.

**Figure S4.**
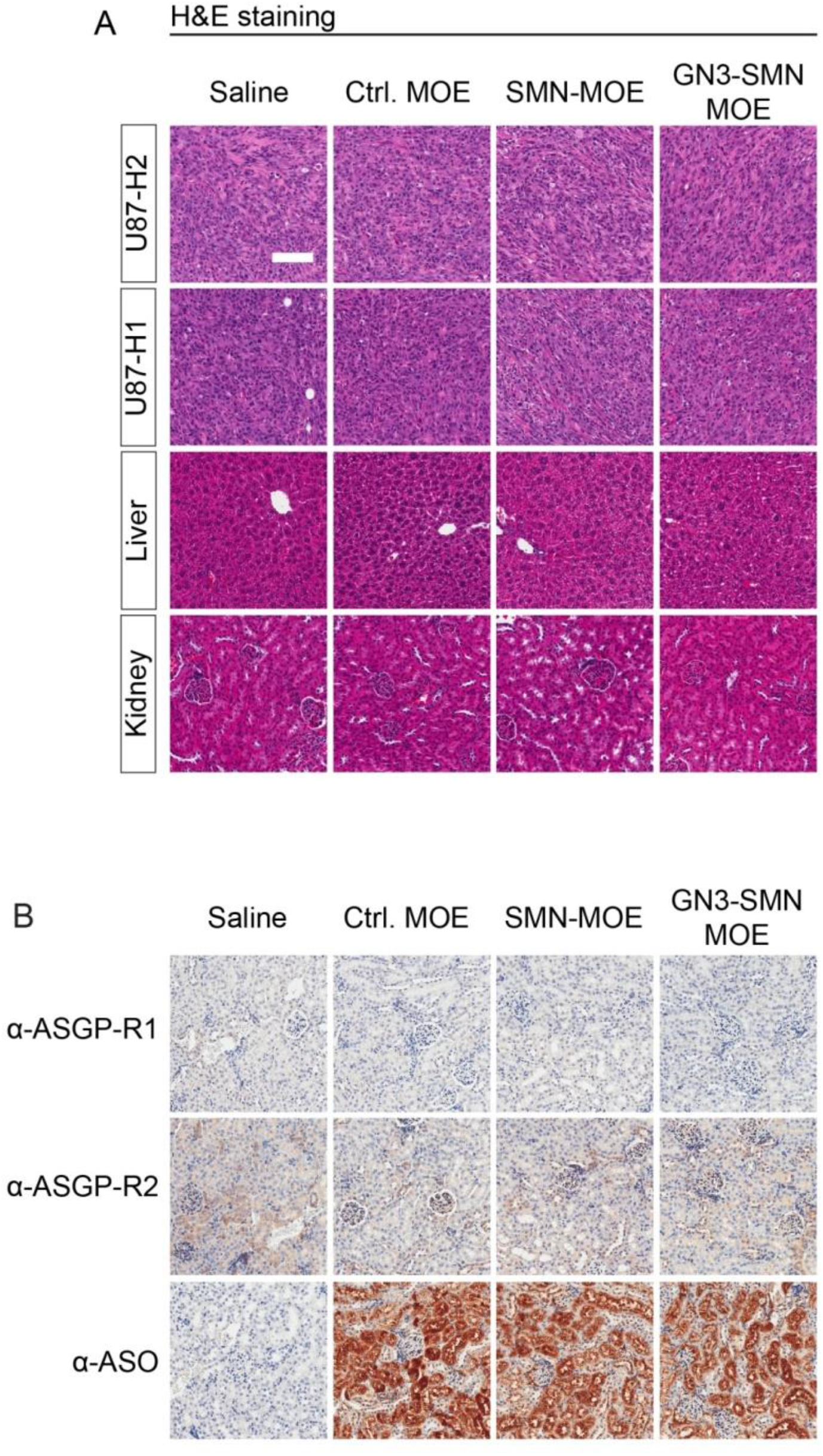
Histological assessment of ASO-treated animals, and ASO distribution in kidney. (A) H&E staining of tumor, liver, and kidney tissue did not reveal any histological abnormalities. (B) Kidney sections from mice treated with saline, unconjugated and GN3-conjugated ASOs stained for ASGP-R1 (top), ASGP-R2 (middle), and ASO (bottom). ASGPR staining was not detected, apart from a faint background signal. ASOs localized in proximal convoluted tubules.

